# Convolutional Sparse Coded Dynamic Brain Functional Connectivity

**DOI:** 10.1101/476663

**Authors:** Jin Yan, Yingying Zhu

## Abstract

Functional brain network has been widely studied in many previous work for brain disorder diagnosis and brain network analysis. However, most previous work focus on static dynamic brain network research. Lots of recent work reveals that the brain shows dynamic activity even in resting state. Such dynamic brain functional connectivity reveals discriminative patterns for identifying many brain disorders. Current sliding window based dynamic brain connectivity framework are not easy to be applied to real clinical applications due to many issues: First, how to set up the optimal sliding window size and how to determine the threshold for the brain connectivity patterns. Secondly, how to represent the high dimensional dynamic brain connectivity pattern in a low dimensional representations for diagnosis purpose. Last, how to deal with the different length dynamic brain network patterns especially when the raw data are of different length. In order to address all those above issues, we proposed a new framework, which employs multiple scale sliding windows and automatically learns a sparse and low ran dynamic brain functional connectivity patterns from raw fMRI data. Furthermore, we are able to measure different length dynamic brain functional connectivity patterns in an equal space by learning a sparse coded convolutional filters. We have evaluated our method with state of the art dynamic brain network methods and the results demonstrated the strong potential of our methods for brain disorder diagnosis in real clinical applications.

## 1 Introduction

Functional magnetic resonance imaging (fMRI) provides a non-invasive way to examine human brain activity. This imaging technique is often referred to as blood oxygenation level dependent (BOLD) imaging [1] because it measures changes of cerebral blood oxygenation that are closely related to neuronal activity [2]. Traditionally fMRI has been used to examine brain activation patterns in health [3–5] and disease [6–8] during the performance of a cognitive or motor task. However, recent studies have begun to use resting-state fMRI (rs-fMRI) to measure regional interactions that occur when a subject is not performing an explicit task [9, 10]. In resting state, fluctuations in spontaneous neural activity are thought to underlie the spontaneous BOLD signal fluctuations. Synchrony, or correlation, between the fluctuations among regions are used to assess inter-regional functional connectivity (FC) in human brain [11, 12].

Many works have been done to extract the functional brain networks from fMRI data. Those works can be divided into two folds: static brain networks and dynamic brain networks. In this work, we focused on the dynamic brain networks study and its application on brain disorder diagnosis. Most of current functional brain network studies calculate the Pearson’s correlation to measure the strength of FC between two brain regions [12, 13]. The dynamic FC patterns are mostly calculated using sliding window techniques [14]. However, those methods suffers from many issues for real applications, such as sliding window size setting up issues, how to reduce high dimensional dynamic brain FC patterns to a low dimensional representations, how to deal with different length fMRI. In order to address those above issues, this work focus on learning a compact representation for dynamic brain networks from different length fMRI data.

In order to achieve this goal, we make several contributions to learn dynamic functional brain network patterns from fMRI data. First, we propose using a one-step multiple scale sliding window to extract the raw dynamic FC patterns from fMRI data. In order to extract robust dynamic FC patterns, we constrain that the learn FC patterns are sparse since the brain connections are biologically sparse. We also constraint that the temporal dynamic changes are a low rank time space since the brain connection changes should always be slow and smooth. Secondly, in order to learn a compact representation for different length dynamic FC patterns, we borrow the idea of convolutional filters from signal processing. As shown in Fig. 1. (a), different length time series could be represented as equal length frequency component in Fourier domain. We proposed to learn a sparse coded filters from the data and project the high dimensional dynamic FC patterns on to the learned filters as shown in Fig. 1. (b). Using this technique, we are able to compare different length dynamic FC patterns extracted from different dataset in the same field. In order to show the performance of our proposed method, we have applied the learned dynamic brain FCs to identify Autism using different training and testing dataset. We compared the performance of our method with state of art work. Experiments results shows that our method has strong potential in dynamic brain network based diagnosis in real clinical applications. The following of this paper are organized as: We first introduce the methods of our work in Section 2. Then, we show the experiment evaluation in Section 3. Section 4 concludes our work and discuss some future directions.

**Fig. 1.**
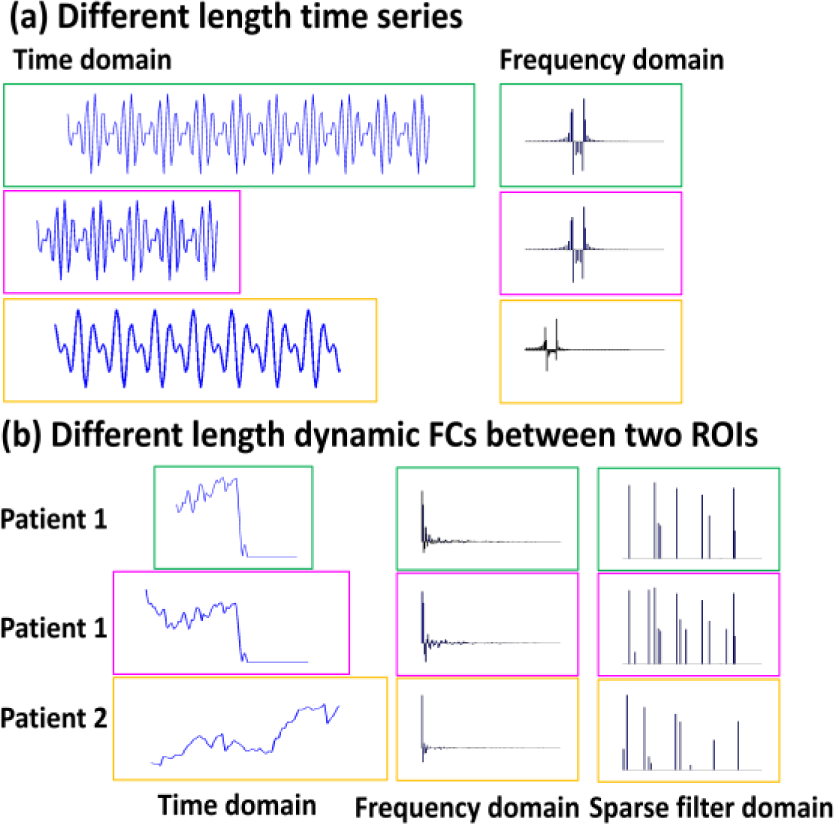
(a) Comparing different length time series using frequency domain information. (b) Comparing the dynamic connection changes with different length in frequency domain and our learned sparse filter domain.

## 2 Methods

We attempt to achieve two goals together in this work: First, learn a clean dynamic FC patterns from the noisy fMRI time series using a set of overlapped multiple-scale sliding windows; Secondly, learn a set of sparse coded convolutional filters to code the learned high dimensional dynamic FC patterns of different length to be equal size.

### 2.1 Robust Dynamic Functional Connectivity

We first describe how we learn the dynamic FC patterns from the original time series in this section. Given a sliding window of size 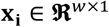 denote the mean BOLD signal calculated in brain region *O_i_,*(*i* = 1, … *,N*), where *w* is the length of time course within the sliding window and *N* is the total number of brain regions under consideration. Conventionally, a *N* × *N* connectivity matrix **S** is used to measure the FC in the whole brain, where each element *S_ij_* quantitatively measures the strength of FC between region *O_i_* and *O_j_* (*i* ≠ *j*). Particularly, the strength of functional connectivity *s_ij_* is assumed to be measurable based on Pearson’s correlation *c*(*x_i_,x_j_*) between observed BOLD signals **x***_i_* and **x***_j_*, where big value of Pearson’s correlation indicates strong functional connectivity. Thresholding on Pearson’s correlation values is commonly used to remove the spurious connection. However, it is not easy to find a good threshold that works for all subjects.

Since fMRI is just an indirect reflection of brain activity, it is difficult to accurately quantify the FC strength only based on signal correlation. To address this issue, we follow the previous work which optimize the reasonable functional connectivities, which should (1) be in consensus with the Pearson’s correlation of low level signals between **x***_i_* and **x***_j_*; (2) use the high level information such as module-to-module connection [15] to guide the measurement of low level region-to-region connectivity strength; and (3) represent sparsity since the brain network is intrinsically efficient to have sparse connections [16]. For convenience, we use 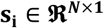 denote i-th column in connectivity matrix **S**, which characters the connections of region *O_i_* with respect to other brain regions. Also, we arrange all Pearson’s correlation values into a *N* × *N* matrix ***C**_i_* = {***c**_ij_|j* = 1, …, *N*} Instead of calculating the connectivity *s_ij_* just based on Pearson’s correlation *c*(**x***i,***x***j*) between observed BOLD signals **x***i* and **x***j*, we optimize the connectivity matrix **S** by integrating the above three criteria:

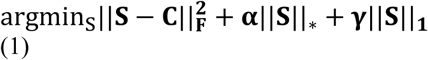

where *α* and *γ* are the scalars which balance the strength of the low rank constraint [17] on **S** (the second term) and the *l*_1_ sparsity constraint [18]on **S** (the third term).

We extend the learning-based FC optimization method to the temporal domain, in order to capture dynamics of functional connectivity. First, we follow the sliding window technique to obtain *T* overlapped multiple scale sliding windows which cover the whole time course for one subject. Let **S***^t^* denote for the FC matrix in sliding window *t*. Then we stack all **S***^t^* along time and form a tensor 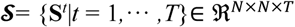 which represents the complete information of dynamic connectivity for each subject. Similarly, we can also construct another tensor 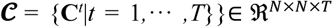, where each **C***^t^_i,j_* = {*c^t^* |*i,j* = 1, … *,N*} is the *N*×*N* matrices in *t-th* sliding window. Next, we propose a learning-based optimization method to characterize dFC using tensor analysis by:

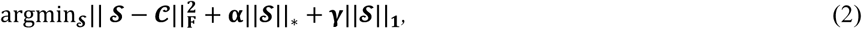

Compared to the objective function in Eq. 1, the objective function here also encourages low rank on the brain connectivity patterns. Since brain in resting state generally transverses a small number of discrete stages during a short period of time, it is reasonable to apply low rank constraint on 𝓢 (by minimizing nuclear norm ||𝓢||∗) to penalize too rapid FC change in the temporal domain and also find the optimal connectivity patterns in each sliding window.

#### Discussion

This above method is able to learn a robust dynamic FC patterns from noisy fMRI time series. However, there are several issues for using these learned dynamic FC patterns for further research or clinical applications. First, this learn dynamic FC pattern is highly redundant and also of high dimension. Secondly, in real applications different patient may have different length fMRI time series, therefore, the learned dynamic FC pattern for each patient varies from each other. If we want to compare them or use them for disease identification, a unified representation or equal size representation should be provided.

### 2.2 Sparse Convolutional Sparse Coded Filters for Dynamic FCs

Motivated by the above two major issues in the learned dynamic FC patterns, we proposed a set of novel sparse coded convolutional dynamic filters for representing redundant dynamic FC patterns compactly with equal size representations. Our method includes two parts: we first learn an over-complete convolutional sparse coded dynamic filters as the dictionary for representing different dynamic FC patterns using training data; then we code each new testing subjects using those learned filters, thus, the different length dynamic FC patterns can be represented as the response of those learned filters, which lead to equal size representation finally. Suppose we have M training patients with fMRI times series data, denote the 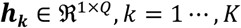, as a convolutional filter, 𝑄 (𝑄 ≪ 𝑇) is the length of this filter, 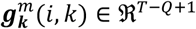 as the response of this filter for ROI *i, j* dynamic connections from subject *m*, 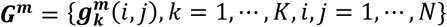 as feature representation for subject **m**, *y_m_* is the clinical label for subject m, we learn an over-complete set of *K* filters using the following objective function,

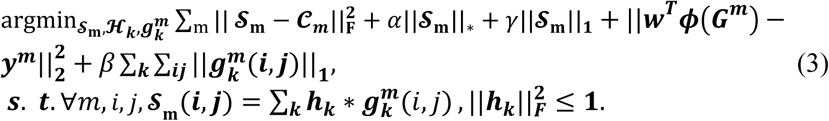

#### Convolutional Feature Coding Testing Subject

After a set of sparse coded filters are learned, for a new testing subject 𝑡𝑒, we jointly learns the robust dynamic FC patterns 𝓢_t_ and the filter response 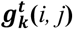 as,

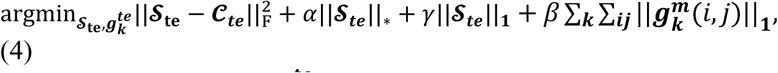

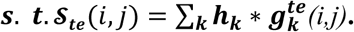

The learned filter response 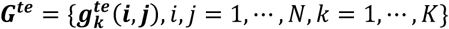 is used as the compact representation for the dynamic FC pattern 𝓢𝐭𝐞.

#### Discussion

Our proposed combined convolutional filters learning and robust dynamic FC pattern learning method has several major improvements from Eq. (2). Firstly, our method automatically learn an over-complete sparse convolutional filters and use the filter response as the compact representation for each patient’s dynamic FC patterns. Secondly, by applying these learned convolutional filters, we are able to represent different length dynamic FC patterns in the same size. Last, those learned convolutional filters can be used for brain functional connectivity pattern related research, such as finding the important biomarkers for brain related diseases.

### 2.3 Optimization

In this section, we will briefly describe how we solve Eq. (4). Eq. (2) can be solved using sub-gradient approach for nuclear norm and 𝑙1 norm optimization. The first part of Eq. (4) can be solve similarity. The convolutional part in Eq. (4) is very hard to be solved directly in the time domain due to the computation cost of convolution. However, it can be easily solved in the Fourier domain since that the convolution in time domain can be represented as multiplication in Fourier domain. Denote *fft* as the Fourier transform, we proposed the new objective function in Fourier domain as,

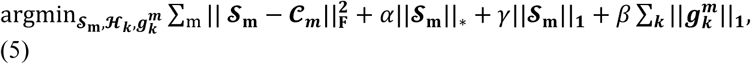

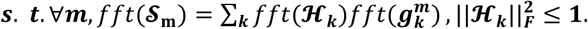

Eq. (5) can be solved iteratively with respect to each parameter iteratively until converge. When we solve the convolutional filter 𝓗𝒌 and filter response 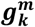, we will convert it in to FFT domain, after the conversion, the convolution can be replaced as multiplication. After we computed the convolution, we will convert it back to time domain for the sparse and low rank constraints. We omit the detailed solutions here due to page limitation. This solution is not the major contribution of our work and we encourage the reviewers to read more details on references [19].

## 3 Experiments

In this section, we evaluate our proposed tensor connectome model of dynamic functional connectivity by comparing the discriminative power in identifying ASD subjects with respect to conventional state-of-the-art methods.

### Subject information

We conducted various experiments on resting-state fMRI images using two Autism data sets in order to demonstrate the generality of our method. We use the Autism Brain Imaging Data Exchange (ABIDE) database including both the data from University of Utah (UU) and University of California Davis (UCD) site. Specifically, 50 NC and 50 ASD subjects are selected from the UU site. 70 NC and 65 ASD subjects are selected from UCD site.

### Data preprocessing

The subjects in our experiments were scanned for six and ten minutes during resting state, respectively, producing 180 time points and 300 time points at a repetition time (TR) of 2s. We processed all these data using Data Processing Assistant for 0 the AAL template with 116 ROIs to the subject image domain and compute the mean BOLD signal in each ROI, where conventional method calculate the 116 × 116 connectivity matrix **S** based on the Pearson’s correlation of mean BOLD signals between any pair of two distinct brain regions.

### Evaluation measurements

We use several quantitative measurements to evaluate not only the classification performance. Besides the widely used Accuracy (ACC) and Accuracy Under ROC Curve (AUC). PCA represents Pearson’s correlation based dynamic FC and feature coded using PCA. OURS represents the learned dynamic FC patterns by our convolutional sparse coded method.

### Experiment setup

Ten-fold cross validation strategy is used in all following experiments. Specifically, we randomly partition all subjects into 10 non-overlapping approximately equal size sets. Then, we use one fold for testing and the remaining folds are used for training. The training subjects are further divided into 5 subsets for another 5-fold inner cross validation, where 4 folds are used as training subset and the last fold is used as the validation subset. The over-completed sparse coded filters are learned from the subjects in the training subset. The low dimension connectome features representations of those training subjects, derived from the robust dynamic FC pattern are further used to train the classic SVM (Support Vector Machine) for classification. The optimal parameters are determined based on validation subset. For each testing subject, we use the approach summarized in Eq. (4) to estimate the dynamic FC feature representations, which is considered as the connectome signature to identify the clinical label of the underlying testing subject. For the competing methods, we apply our multiple scale sliding window strategy to calculate the dynamic functional connectivity feature representation (based on Pearson’s correlation and optimal thresholding on validation dataset) for each subject. We first manually set up the sliding window size which ranges from 20 to 100 of the entire time course. Please note that the brain connection pattern are unstable is the window size is smaller than 20. In optimizing the dynamic FC pattern, we set the shift of sliding window to 1 TR, in order to fully capture the dynamics of FC. In order to reduce feature dimension, we follow the work in to use classic PCA (Principle Component Analysis) model [20]to encode the low dimensional connectome feature representation for each subject for comparison.

### Evaluation of learned dynamic FC patterns in NC/ASD classification using the same dataset

We first evaluate the performance of our model using the same dataset for training and testing. 10-fold cross-validation strategy is employed here on UU and UCD dataset. The NC/ASD classification results using different sliding window setup on UU and UCD dataset are shown in Fig. 2 (a) and (b) respectively. It is shown that, first, the optimal sliding window size is 40 for two datasets; secondly, multiple scale sliding window setup achieves the best performance for all methods on two datasets, which improves *>* 2% in terms of Accuracy compared to best performance achieved by fixed sliding window size; our learned 4D tensor feature representation improves the performance at least 4.5% in terms of Accuracy compared with the conventional Pearson’s correlation method using multiple scale sliding windows.

**Fig. 2.**
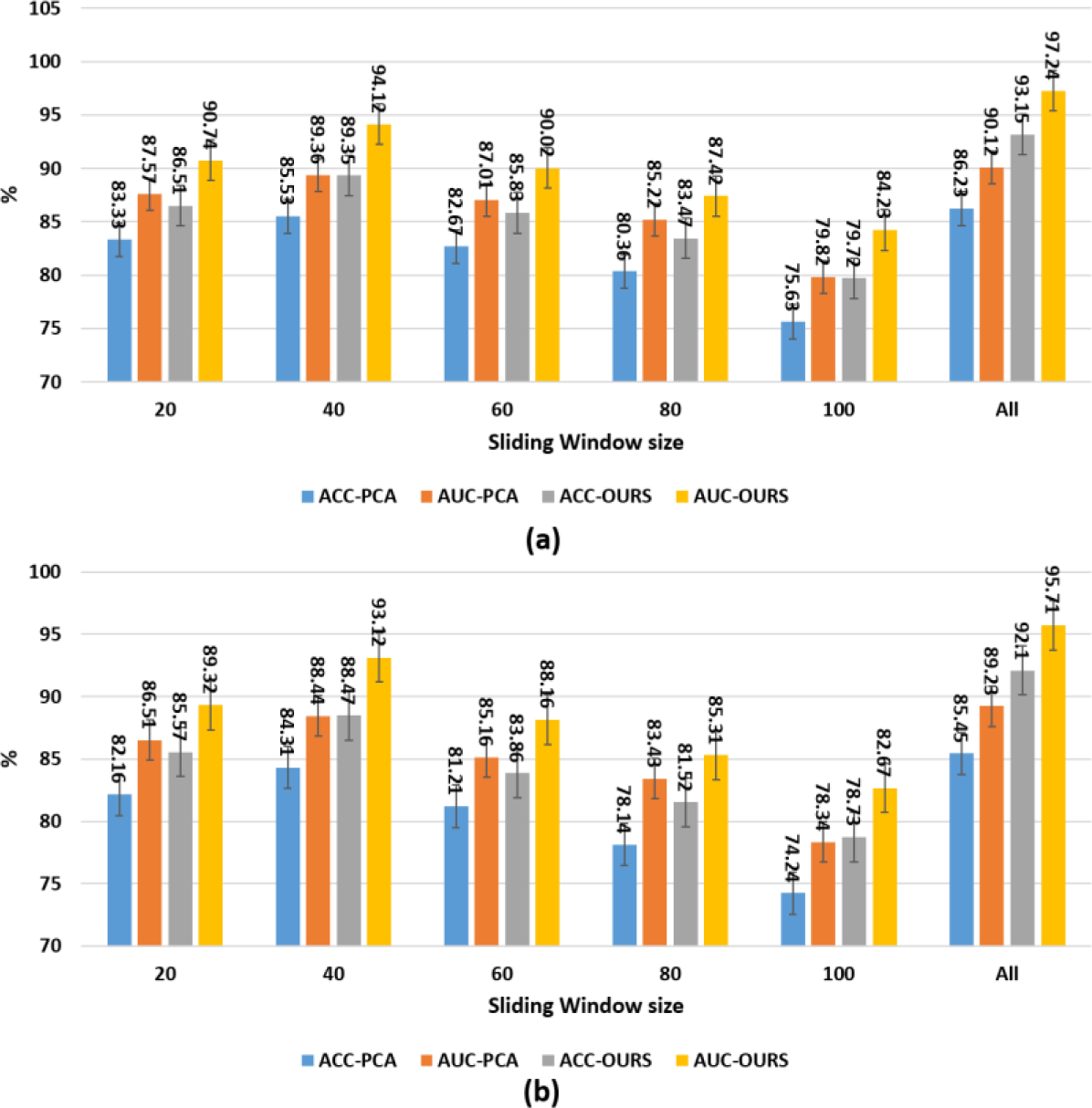
Evaluation of learned dynamic FC patterns using the same training and testing dataset. (a) Identifying ASD subjects on UU dataset w.r.t. sliding window size. (b) Identifying ASD subjects on UCD dataset w.r.t. sliding window size.

### Evaluation of learned dynamic FC patterns in NC/ASD classification using different dataset

To evaluate the generalization of the learned dynamic FC pattern representations, we select the training data and testing data from different sites. Two experiments are conducted here: first, we split the UU and UCD data into ten non-overlapped folds and use 9 folds from UU as the training data and one fold from UU as the testing data; then, we switch the training and testing dataset. Fig. 3 shows the performance of conventional Pearson’s correlation patterns coded by PCA and our learned dynamic FC patterns with respect to different sliding window sizes. The performance is sensitive to different window size setting up. All competing methods achieve the best performance when the sliding window is 40 and multiple scale sliding window setting up improves the performance about *>* 1% compared to sliding window size 40 for all competing methods. Compared with the conventional Pearson’s correlation patterns coded by PCA [21], our learned dynamic FC pattern improves the ASD classification performance *>* 4% on Accuracy using multiple scale sliding windows.

**Fig. 3.**
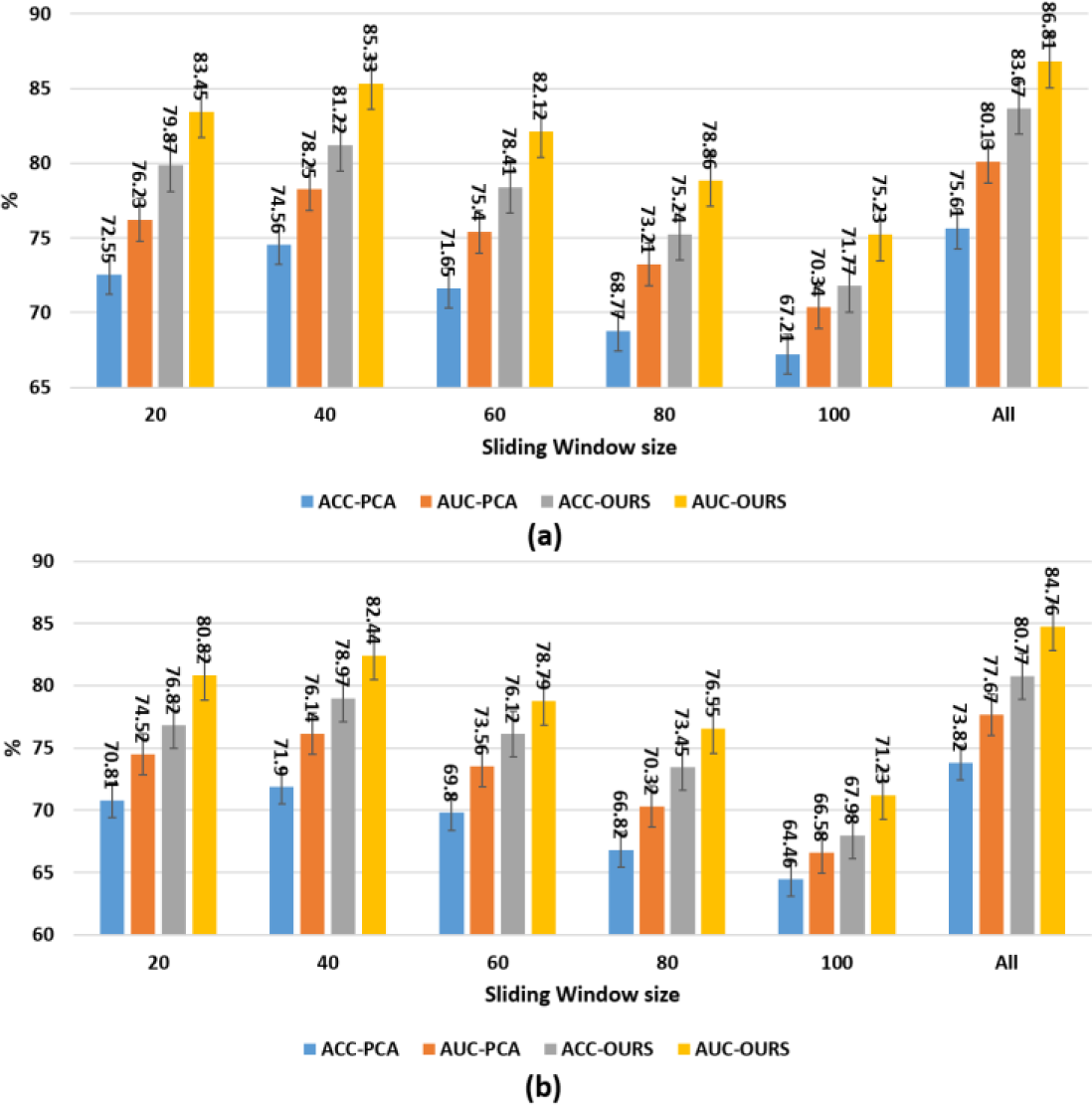
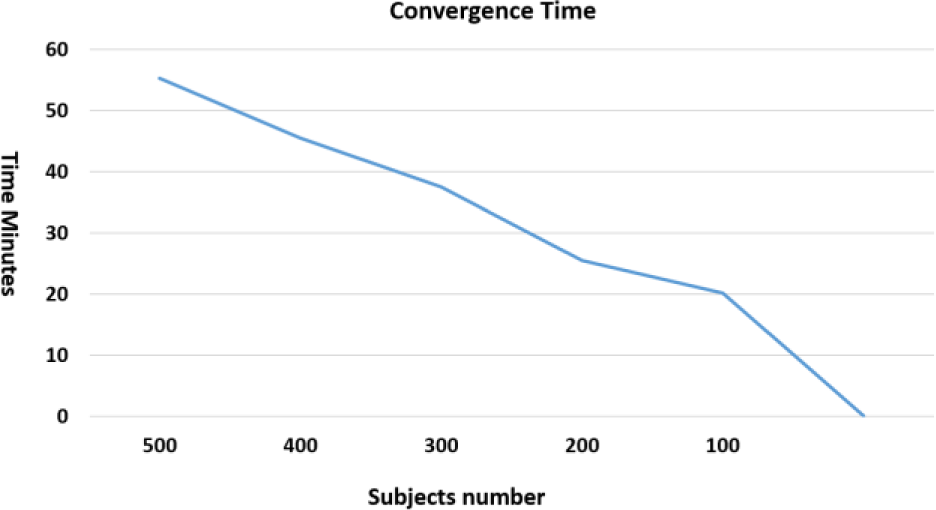
Evaluation of learned dynamic FC patterns using different training and testing dataset. (a) Identifying ASD subjects on UU dataset w.r.t. sliding window size trained using UCD dataset. (b) Identifying ASD subjects on UCD dataset w.r.t. sliding window size trained using UU dataset. The computation time of our method on different data size. As we can see, the computational cost increase linearly with the size of data.

### Convergence of our method

In order to show the convergence of our method. We select different size training data set. The training data size varies from 50 to 500. We show the computation time cost in **Fig. 4.** One can see that the computation cost increases linearly with the number of subjects. We expect that our method can handle large size data well. And since the most time consuming process in medical imaging based diagnosis is the data pro-processing, such as registration and segmentation, we believe that our method can be use in practical problems.

**Fig. 4.**
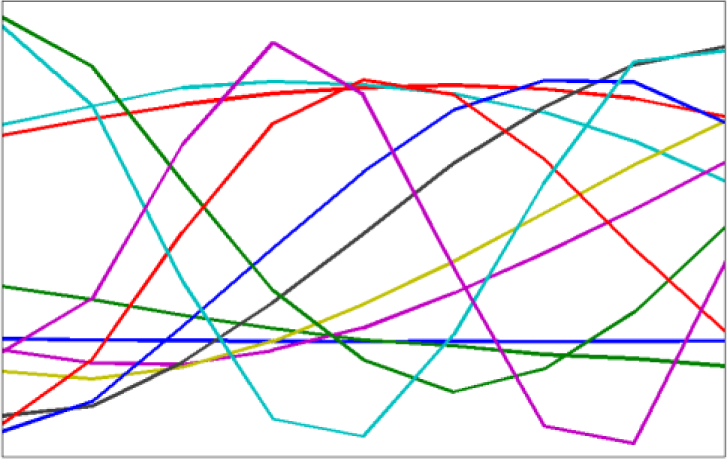
Visualization of learned filters by our method.

### Visualization of learned convolutional filters

We show the top 12 learned convolutional filters learned by our methods in **Fig. 4.** It is very smooth and have different slopes. This results suggest that the brain connectivity changes are smooth and periodically.

## 4 Conclusion

In this work, we aims at solving the dynamic brain functional network extraction problem. We proposed a multiple scale sliding window framework combined with low rank sparse constraint to learn a robust dynamic brain functional network patterns from fMRI data. In order to compare different length dynamic brain functional network patterns, we further proposed to learn a sparse convolutional coded filters for representing different length dynamic brain networks. We have evaluated our method using different dataset of different length fMRI data. Promising results shows the potential of our method for real clinical applications. Future work will explore the application of the dynamic brain network model for more brain disorder diagnosis and brain network analysis.

## 5 Conflict of Interest

The authors whose names are listed immediately below certify that they have NO affiliations with or involvement in any organization or entity with any financial interest (such as honoraria; educational grants; participation in speakers’ bureaus; membership, employment, consultancies, stock ownership, or other equity interest; and expert testimony or patent-licensing arrangements), or non-financial interest (such as personal or professional relationships, affiliations, knowledge or beliefs) in the subject matter or materials discussed in this manuscript. Author names: The authors whose names are listed immediately below report the following details of affiliation or involvement in an organization or entity with a financial or non-financial interest in the subject matter or materials discussed in this manuscript. Please specify the nature of the conflict on a separate sheet of paper if the space below is inadequate.

